# Genome sequence of the banana aphid, *Pentalonia nigronervosa* Coquerel (Hemiptera: Aphididae) and its symbionts

**DOI:** 10.1101/2020.04.25.060517

**Authors:** Thomas C. Mathers, Sam T. Mugford, Saskia A. Hogenhout, Leena Tripathi

**Affiliations:** Department of Crop Genetics, John Innes Centre, Norwich Research Park, Norwich, United Kingdom; International Institute of Tropical Agriculture (IITA), P.O. Box 30709-00100, Nairobi, Kenya

**Keywords:** Hemiptera, genome assembly, insect vector, plant pest, phylogenomics

## Abstract

The banana aphid, *Pentalonia nigronervosa* Coquerel (Hemiptera: Aphididae), is a major pest of cultivated bananas (*Musa* spp., order Zingiberales), primarily due to its role as a vector of *Banana bunchy top virus* (BBTV), the most severe viral disease of banana worldwide. Here, we generated a highly complete genome assembly of *P. nigronervos*a using a single PCR-free Illumina sequencing library. Using the same sequence data, we also generated complete genome assemblies of the *P. nigronervos*a symbiotic bacteria *Buchnera aphidicola* and *Wolbachia*. To improve our initial assembly of *P. nigronervos*a we developed a k-mer based deduplication pipeline to remove genomic scaffolds derived from the assembly of haplotigs (allelic variants assembled as separate scaffolds). To demonstrate the usefulness of this pipeline, we applied it to the recently generated assembly of the aphid *Myzus cerasi*, reducing the duplication of conserved BUSCO genes by 25%. Phylogenomic analysis of *P. nigronervos*a, our improved *M. cerasi* assembly, and seven previously published aphid genomes, spanning three aphid tribes and two subfamilies, reveals that *P. nigronervos*a falls within the tribe Macrosiphini, but is an outgroup to other Macrosiphini sequenced so far. As such, the genomic resources reported here will be useful for understanding both the evolution of Macrosphini and for the study of *P. nigronervos*a. Furthermore, our approach using low cost, high-quality, Illumina short-reads to generate complete genome assemblies of understudied aphid species will help to fill in genomic black spots in the diverse aphid tree of life.

## Introduction

Aphids are economically important plant pests that cause damage to crops and ornamental plant species through parasitic feeding on plant sap and via the transmission of plant viruses (Van Emden and Harrington 2017). Of approximately 5,000 aphid species, around 100 have been identified as significant agricultural pests. Despite their economic importance, little to no genomic resources exist for many of these species or their relatives, hindering efforts to understand the evolution and ecology of aphid pests. To date, genome sequencing efforts have focused on members of the aphid tribe Macrosiphini (within subfamily Aphidinae), including the widely studied aphids *Acyrthosiphon pisum* (pea aphid) (IAGC 2010; Li *et al.* 2019; Mathers *et al.* 2020) and *Myzus persicae* (green peach aphid) (Mathers *et al.* 2017, 2020), as well as other important pest species such as *Diuraphis noxia* (Russian wheat aphid) (Nicholson *et al.* 2015). Recently, additional genome sequences have become available for members of the tribe Aphidini (also in the subfamily Aphidinae) (Wenger *et al.* 2017; Thorpe *et al.* 2018; Chen *et al.* 2019; Quan *et al.* 2019; Mathers 2020) and the subfamily Lanchiniae (Julca *et al.* 2019), broadening the phylogenetic scope of aphid genomic resources. However, many clades of the aphid phylogeny are still missing or underrepresented in genomic studies.

The banana aphid, *Pentalonia nigronervosa* Coquerel (Hemiptera: Aphididae), is a major pest of cultivated bananas (*Musa* spp., order Zingiberales) and is widely distributed in tropical and subtropical regions where bananas are grown (Waterhouse 1987). Like other aphid species, *P. nigronervosa* feeds predominantly from the phloem of its plant host. Intensive feeding can kill or affect the growth of young banana plants. However, direct feeding damage to adult plants is often negligible. Instead, the banana aphid causes most economic damage as a vector of plant viruses, some of which induce severe disease symptoms and substantial yield loss of banana (Dale 1987; Sharman *et al.* 2008; Savory and Ramakrishnan 2015). In particular, *P. nigronervosa* is the primary vector of the *Banana bunchy top virus* (BBTV), the most severe viral disease of banana worldwide (Dale 1987).

*P. nigronervosa* carries at least two bacterial symbionts: *Buchnera aphidicola* and *Wolbachia* (de Clerck *et al.* 2014). *Buchnera aphidicola* is an obligate (primary) symbiont present in almost all aphid species and provides essential amino acids to the aphids (Baumann 1995; Douglas 1998; Hansen and Moran 2011; Shigenobu and Wilson 2011). In contrast, *Wolbachia* is considered a facultative (secondary) symbiont and is found in a few aphid species at low abundance (Augustinos *et al.* 2011; Jones *et al.* 2011). Interestingly, *Wolbachia* is found systematically across the *P. nigronervosa* range (de Clerck *et al.* 2014) and is also present in the closely related species *P. caladii* van der Goot (Jones *et al.* 2011), which rarely colonizes banana, and prefers other plant species of the order Zingiberales (Foottit *et al.* 2010). Possibly, *Wolbachia* provides essential nutrients and vitamins to the *Pentalonia* spp or/and protects them from plant-produced defense molecules such as anti-oxidants or phenolic compounds of banana (Hosokawa *et al.* 2010).

Here, we generate highly complete genome assemblies of *P. nigronervosa* and its symbiotic bacteria *Buchnera aphidicola* and *Wolbachia,* using a single PCR-free Illumina sequencing library. Phylogenomic analysis reveals that *P. nigronervos*a falls within the aphid tribe Macrosiphini, but is an outgroup to other Macrosiphini sequenced so far. As such, the genomic resources reported here will useful for understanding the evolution of Macrosphini, and for the study of *P. nigronervosa*.

## Methods

### Aphid rearing and sequencing library construction

A lab colony of *P. nigronervosa* was established from a single asexually reproducing female collected initially from the IITA’s banana field at the International Livestock Research Institute (ILRI) Nairobi, Kenya. A single colony of *P. nigronervosa* was collected from a field-grown banana plant and introduced on an eight-week-old potted tissue culture banana plant in an insect-proof cage, placed in a glasshouse under room temperature and natural light. Pure aphid colonies were propagated by transferring a single aphid from the potted banana plant to another fresh young banana plant in the glasshouse every eight weeks. Aphids from this colony were used for all subsequent DNA and RNA extractions. Genomic DNA was extracted from a single individual with a modified CTAB protocol (based on Marzachi *et al.* 1998) and sent to Novogene (China), for library preparation and sequencing. Novogene prepared a PCR free Illumina sequencing library using the NEBNext Ultra II DNA Library Prep Kit for Illumina (New England Biolabs, USA), with the manufacturers protocol modified to give a 500 bp – 1 kb insert size. This library was sequenced on an Illumina HiSeq 2500 instrument with 250 bp paired-end chemistry. To aid scaffolding and genome annotation, we also generated a high coverage, strand-specific, RNA-seq library. RNA was extracted from whole bodies of 20-25 individuals using Trizol (Signma) followed by clean-up and on-column DNAse digestion using RNeasy (Qiagen) according to the manufactures’ protocols, and sent to Novogene (China) where a sequencing library was prepared using the NEBNext Ultra Directional RNA Library Prep Kit for Illumina (New England Biolabs, USA). This library was sequenced on an Illumina platform with 150 bp paired-end chemistry.

### De novo *genome assembly and quality control*

Raw sequencing reads were processed with trim_galore (http://www.bioinformatics.babraham.ac.uk/projects/trim_galore) to remove adapters and then assembled using Discovar *de novo* (https://software.broadinstitute.org/software/discovar/blog/) with default parameters. The content of this initial assembly was assessed with Benchmarking Universal Single-Copy Orthologs (BUSCO) v3.0 (Simão *et al.* 2015; Waterhouse *et al.* 2018) using the Arthropoda gene set (n = 1,066) and by k-mer analysis with the k-mer Analysis Toolkit (KAT) v2.2.0 (Mapleson *et al.* 2017), comparing k-mers present in the raw sequencing reads to k-mers found in the genome assembly with KAT comp. We identified a small amount of k-mer content that was present twice in the genome assembly but that had k-mer coverage in the reads of a single-copy region of the genome, indicating the assembly of haplotigs (allelic variants that are assembled into separate contigs) (**Supplementary Figure 1a**). To generate a close-to-haploid representation of the genome, we applied a strict filtering pipeline to the draft assembly based on k-mer analysis and whole genome self-alignment. Firstly, the k-mer coverage of the homozygous portion of the genome was estimated with KAT distanalysis, which decomposes the k-mer spectra generated by KAT comp into discrete distributions corresponding to the number of times their content is found in the genome. Then, for each scaffold in the draft assembly, we used KAT sect to calculate the median k-mer coverage in the reads and the median k-mer coverage in the assembly. Scaffolds that had medium k-mer coverage of 2 in the assembly and median k-mer coverage in the reads that fell between the upper and lower bounds of homozygous genome content (identified by KAT distanalysis), were flagged as putative haplotigs. We then carried out whole genome self-alignment with nucmer v4.0.0beta2 (Marçais *et al.* 2018) and removed putative haplotigs that aligned to another longer scaffold in the genome with at least 75% identity and 25% coverage. The deduplicated assembly was then checked again with BUSCO and KAT comp to ensure that no (or minimal) genuine homozygous content had been lost from the assembly.

The deduplicated draft assembly was screened for contamination based on manual inspection of taxon-annotated GC content coverage plots (“blobplots”) generated with BlobTools v1.0.1 (Kumar *et al.* 2013; Laetsch and Blaxter 2017). Genomic reads were aligned to the deduplicated draft assembly with BWA mem (Li 2013) and used to estimate average coverage per scaffold. Additionally, each scaffold in the assembly was compared to the NCBI nucleotide database (nt) with BLASTN v2.2.31 (Camacho *et al.* 2009). Read mappings and blast results were then passed to BlobTools which was used to create “blobplots” annotated with taxonomy at the order- and genus-level. Using this approach, we were able to identify and remove scaffolds corresponding to bacterial symbionts and scaffolds that had aberrant coverage and GC content patterns that are likely contaminants.

Finally, to further improve contiguity and gene-level completeness, we performed an additional round of scaffolding using our high coverage RNA-seq data with P_RNA_scaffodler (Zhu *et al.* 2018). RNA-seq reads were trimmed for adapters and low-quality bases with trim_galore and aligned to the deduplicated and cleaned assembly with HISAT2 v2.0.5 [-k 3 -pen-noncansplice 1000000] (Kim *et al.* 2015). The resulting BAM file was then passed to P_RNA_scaffolder along with the draft assembly, and scaffolding performed with default settings. Gene-level completeness was assessed before and after RNA-seq scaffolding with BUSCO and final runs of KAT comp and BlobTools were performed to check the quality and completeness of the assembly.

### Genome annotation

Repeats were identified and soft-masked in the frozen genome assembly using RepeatMasker v4.0.7 [-e ncbi -species insecta -a -xsmall -gff] (Smit *et al.* 2005) with the Repbase (Bao *et al.* 2015) Insecta repeat library. We then carried out gene prediction on the soft-masked genome using the BRAKER2 pipeline v2.0.4 (Lomsadze *et al.* 2014; Hoff *et al.* 2015) with RNA-seq evidence. BRAKER2 uses RNA-seq data to create intron hints and train a species-specific Augustus (Stanke *et al.* 2006, 2008) model which is subsequently used to predict protein coding genes, taking RNA-seq evidence into account. RNA-seq reads were aligned to the genome with HISAT2 v2.0.5 [--max-intronlen 25000 --dta-cufflinks --rna-strandness RF] and the resulting BAM file passed to BRAKER2, which was run with default settings. Completeness of the BRAKER2 gene set was assessed using BUSCO with the Arthropoda gene set (n=1,066). We generated a functional annotation of the predicted gene models using InterProScan v5.22.61 (Enright *et al.* 2002; Jones *et al.* 2014).

### *Upgrading* Myzus cerasi *v1.1*

To demonstrate the usefulness of our k-mer based deduplication pipeline, we applied it to the published short-read assembly of *M. cerasi* (Mycer_v1.1) (Thorpe *et al.* 2018). We ran the pipeline as for *P. nigronervosa*, using the PCR-free Illumina reads that were originally used to assemble Mycer_v1.1 (NCBI bioproject PRJEB24287) and scaffolded the deduplicated assembly using RNA-seq data from Thorpe et al. (2016) (PRJEB9912) with P_RNA_scaffolder. RNA-seq reads were first trimmed for low quality bases and adapters with trim_galore, retaining reads where both members of a pair were at least 75 bp long after trimming. The deduplicated, scaffolded, assembly was ordered by size and assigned a numbered scaffold ID to create a frozen release for downstream analysis (Mycer_v1.2). Mycer_v1.2 was then soft-masked with RepeatMasker using the Repbase Insecta repeat library and protein coding genes predicted with BRAKER2 using the Thorpe et al. (2016) RNA-seq.

### Phylogenomic analysis of aphids

Protein sequences from *P. nigronervosa*, our upgraded *M. cerasi* genome, and seven previously published aphid genomes (**Supplementary Table 1**), were clustered into orthogroups with Orthofinder v2.2.3 (Emms and Kelly 2015, 2018). Where genes had multiple annotated transcripts, we used the longest transcript to represent the gene model. Orthofinder is a comparative genomics pipeline that reconstructs orthogroups, estimates the rooted species tree, generates rooted gene trees, and infers orthologs and gene duplication events using the rooted gene trees, providing a rich resource for downstream comparative analysis. We ran Orthofinder in multiple sequence alignment mode [-M msa -S diamond -T fasttree] using MAFFT (Katoh and Standley 2013) to align orthogroups and FastTree (Price *et al.* 2010) to infer Maximum Likelihood gene trees for each orthogroup. The species tree was then estimated based on a concatenated alignment of all conserved single-copy orthogroups and rooted using evidence from gene duplications with STRIDE (Emms and Kelly 2017).

### Data availability

Sequence data for this project has been deposited in the NCBI short read archive under project accession number PRJNA628023. The *P. nigronervosa* genome assembly and annotation, the updated *M. cerasi* genome assembly and annotation, orthogroup clustering results and code to run our assembly de-duplication pipeline are available for download from Zenodo (https://10.5281/zenodo.3765644).

## Results and Discussion

### P. nigronervosa *genome assembly and annotation*

In total we generated 23 Gb of PCR-free Illumina genome sequence data (~61x coverage of the *P. nigronervosa* genome) and 18 Gb of strand-specific RNA-seq data from a clonal lineage of *P. nigronervosa* (**Supplementary Table 2**). Using these data, we generated a *de novo* genome assembly of *P. nigronervosa* (Penig_v1). Penig_v1 is assembled into 18,348 scaffolds totaling 375 Mb of sequence with an N50 of 104 Kb (contig N50 = 64 Kb, n = 20,873; **Table 1**). The assembly is highly complete, with little duplicated or missing content (**Figure 1a**), and has excellent representation of conserved arthropod genes (95% complete and single-copy), meeting or exceeding the completeness of other published aphid genomes (**Figure 1b**). Furthermore, taxon annotated “blob-plots” show that Penig_v1 is free from obvious contamination (**Supplementary Figure 2**). Gene prediction using BRAKER2 with RNA-seq evidence resulted in the annotation of 27,698 protein coding genes and 29,708 transcripts. Completeness of the gene set reflects that of the genome assembly with 93% of BUSCO Arthropoda genes present as complete single copies in the annotation (**Supplementary Figure 3**). We were able to assign functional domains to 12,869 (47 %) of the annotated gene models (**Supplementary Table 3**). Statistics for the final assembly and annotation of *P. nigronervosa* are summarised in **Table 1**.

**Figure 1:**
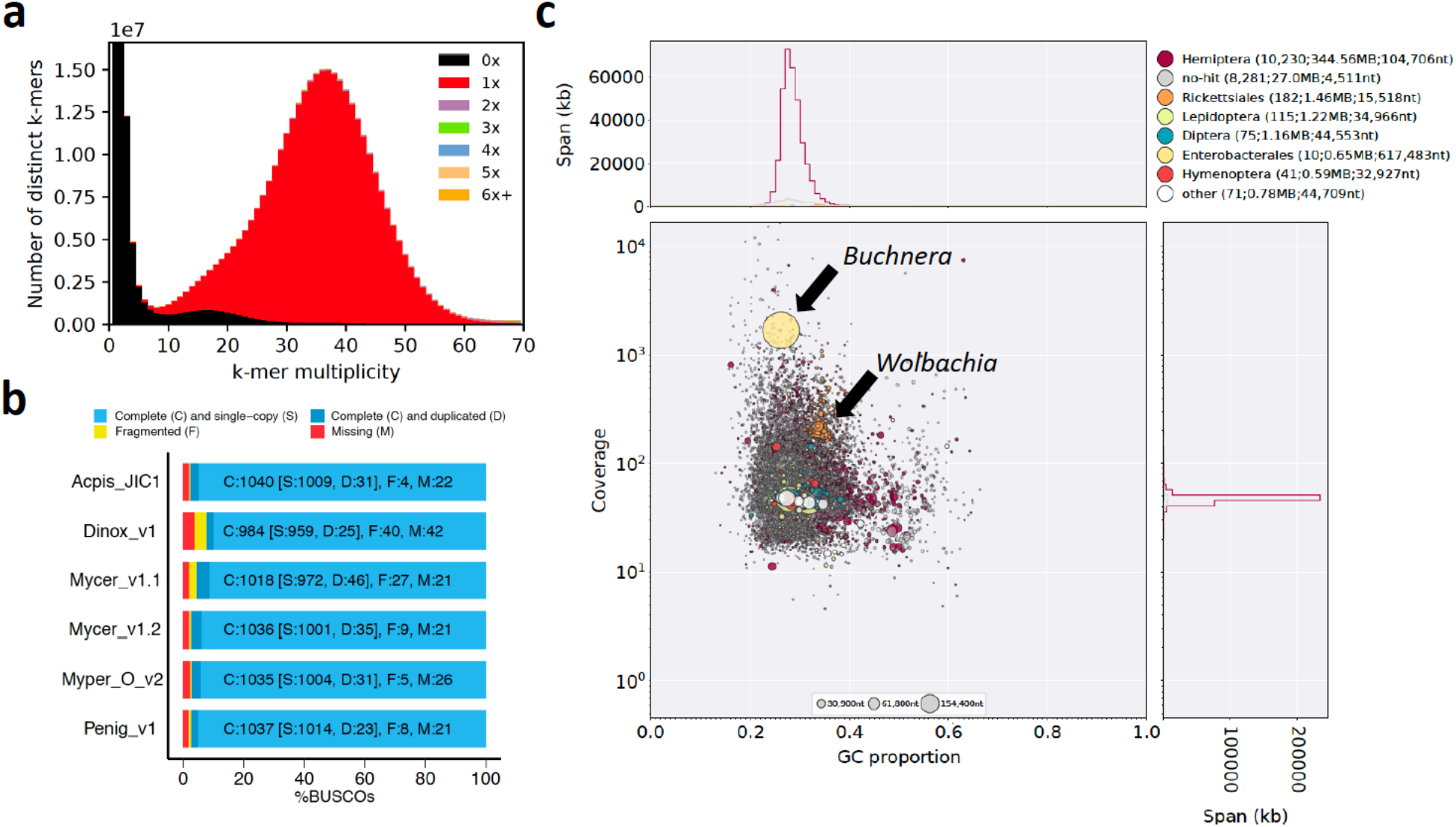
The *P. nigronervosa* genome assembly is complete and free from duplication and contamination. (**a**) KAT k-mer spectra plot comparing k-mer content of PCR-free *P. nigronervosa* Illumina reads to k-mer content of the final *P. nigronervosa* genome assembly (Penig_v1). Colours indicate how many times fixed length words (k-mers) from the reads appear in the assembly. Red indicates k-mers found only once in the assembly, black indicates content present in the reads but missing from the assembly and other colours indicate k-mers that are duplicated in the assembly. The x-axis shows the number of times each k-mer is found in the reads (k-mer multiplicity) and the y-axis shows the count of distinct k-mers in 1x k-mer multiplicity bins. (**b**) BUSCO analysis of Penig_v1, our updated assembly of *M. cerasi* (Mycer_v1.2) and published Macrosiphini genome assemblies. Myper_O_v2 = *Myzus persicae* clone O v2, Acpis_JIC1 = *Acyrthosiphon pisum* clone JIC1, Mycer_v1.1 = *Myzus cerasi* v1.1 and Dnox_v1 = *Diuraphis noxia* v1. The genomes were assessed using the Arthropoda gene set (n=1,066). (**c**) Taxon-annotated GC content-coverage plot of the *P. nigronervosa* Discovar *de novo* genome assembly (post deduplication and prior to RNA-seq scaffolding – see Methods) showing co-assembly of the aphid and its symbionts. Each circle represents a scaffold in the assembly, scaled by length, and coloured by order-level NCBI taxonomy assigned by BlobTools. The X axis corresponds to the average GC content of each scaffold and the Y axis corresponds to the average coverage based on alignment of *P. nigronervosa* PCR-free Illumina short reads. Marginal histograms show cumulative genome content (in Kb) for bins of coverage (Y axis) and GC content (X axis). Arrows highlight scaffolds assigned to the symbiotic bacteria *Buchnera aphidicola* and *Wolbachia* which were removed from the final assembly (**Supplementary Figure 2**).

**Table 1:**
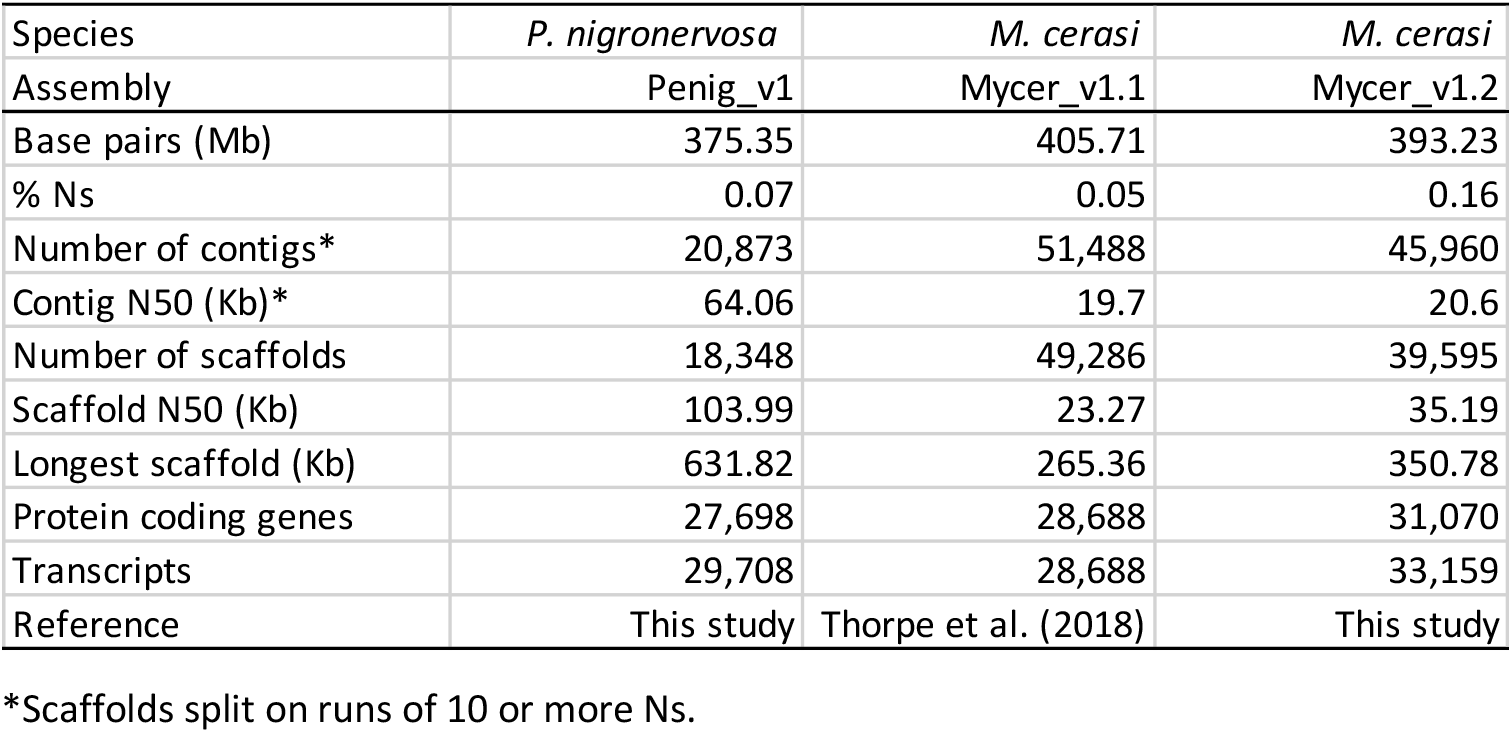
Genome assembly and annotation statistics for *P. nigronervosa* and *M. cerasi*.

*P. nigronervosa* in known to harbour the obligate aphid bacterial endosymbiont *Buchnera aphidicola* and a secondary symbiont, *Wolbachia*, that is found systematically across the species range (de Clerck *et al.* 2014). We identified both symbiotic bacteria in the initial discovar *de novo* assembly of *P. nigronervosa* (**Figure 1c**). *B. aphidicola* BPn was assembled into a single circular scaffold 617 KB in length, along with 2 plasmids. The *Wolbachia* WolPenNig assembly was more fragmented (1.46 Mb total length, 182 scaffolds, N50 = 15.5kb). Despite this, the WolPenNig assembly is likely highly complete as it is similar in size to both a more contiguous long-read assembly of a strain found in the soybean aphid (1.52 Mb total length, 9 contigs, N50 = 841 Kb [Mathers 2020]) and to the reference assembly of *Wolbachia* wRi (Klasson *et al.* 2009) from *Drosophila simulans* (1.44 Mb total length, 1 contig). Furthermore, BUSCO analysis using the proteobacteria gene set (n = 221) reveals that WolPenNig has similar gene-level completeness to these high-quality assemblies, with 81% of BUSCO genes found as complete, single copies (**Supplementary Figure 4**).

### *Upgrading the* Myzus cerasi *genome assembly and annotation*

The initial discovar *de novo* assembly of *P. nigronervosa* was moderately improved by applying our deduplication pipeline and by scaffolding the assembly with RNA-seq data. Compared to the raw discovar *de novo* assembly, contiguity increased by 8% (scaffold N50 = 104 kb vs 96 Kb). Furthermore, the number of fragmented BUSCO Arthropoda genes was reduced from 11 to 8 indicating improved representation of the gene space in the processed assembly. Because the pipeline removes scaffolds that are predominantly made up of erroneously duplicated k-mers, these improvements were achieved without compromising genuine single-copy genome content (**Supplementary Figure 1b**). This approach will likely benefit other low-cost aphid genome assembly projects that use short-read sequencing, particularly when heterozygosity is high. To demonstrate this, we attempted to improve the published genome assembly of *Myzus cerasi* (Mcer_v1.1) (Thorpe *et al.* 2018), using publicly available data. Mcer_v1.1 is made up of 49,286 scaffolds, and k-mer analysis shows high heterozygosity and the presence duplicated content, likely the result of assembling haplotigs (**Supplementary Figure 5a**). We applied our deduplication and RNA-seq scaffolding pipeline to Mcer_v1.1 to create Mcer_v1.2. In total we removed 12.9 Mb of putatively duplicated content from Mcer_v1.1, reducing the assembly size from 405.5 to 392.6 Mb (**Table 1**). The updated assembly is 52% more contiguous than Mcer_v1.1 (scaffold N50 = 35 Kb vs 23 Kb; **Table 1**) and BUSCO analysis indicates that Mcer_v1.2 better represents the gene space, with fewer duplicated (35 vs. 46) and fragmented (9 vs. 27) BUSCO Arthropoda genes (**Figure 1b**). As with Pnig_v1, these improvements were achieved without loss of genuine single-copy genome content (**Supplementary Figure 5b**). We annotated protein coding genes in Mcer_v1.2 with BRAKER2 using RNA-seq evidence, identifying 31,070 protein coding genes with 33,159 transcripts. Again, BUSCO analysis of the updated gene set indicates significant improvement over Mcer_v1.1, with the number of missing and fragmented BUSCO Arthropoda genes reduced from 65 to 20 and 55 to 20 respectively, and overall completeness increased by 8% from 946 to 1,026 BUSCO Arthropoda genes (**Supplementary Figure 3**).

### P. nigronervosa *is an outgroup to other sequenced Macrosiphini*

To investigate the phylogenetic position of *P. nigronervosa* within aphids we carried out orthology clustering of 223,889 protein sequences from *P. nigronervosa*, our improved *M. cerasi* annotation, and seven previously published aphid genomes (Nicholson *et al.* 2015; Thorpe *et al.* 2018; Chen *et al.* 2019; Mathers 2020; Mathers *et al.* 2020). Although the number of aphid species with sequenced genomes is still low, the included species span three aphid tribes (Macrosphini, Aphidini and Lachnini) and approximately 100 million years of aphid evolution (Kim *et al.* 2011; Hardy *et al.* 2015; Julca *et al.* 2019). In total, 204,139 genes (85%) were clustered into 22,759 orthogroups, 4,721 of which are conserved and single-copy in all species (**Supplementary table 4**). Maximum likelihood phylogenetic analysis using a concatenated alignment of the single-copy orthogroups produced a fully resolved species tree with 100% support at all nodes (**Figure 2**). Macrosiphini and Aphidini are recovered as monophyletic groups in agreement with previous analyses based on a small number of genes (von Dohlen *et al.* 2006; Choi *et al.* 2018) and a recent phylogenomic analysis of aphids and other insects (Julca *et al.* 2019). *P. nigronervosa* is placed as an outgroup to other, previously sequenced, members of Macrosiphini (**Figure 2**).

**Figure 2:**
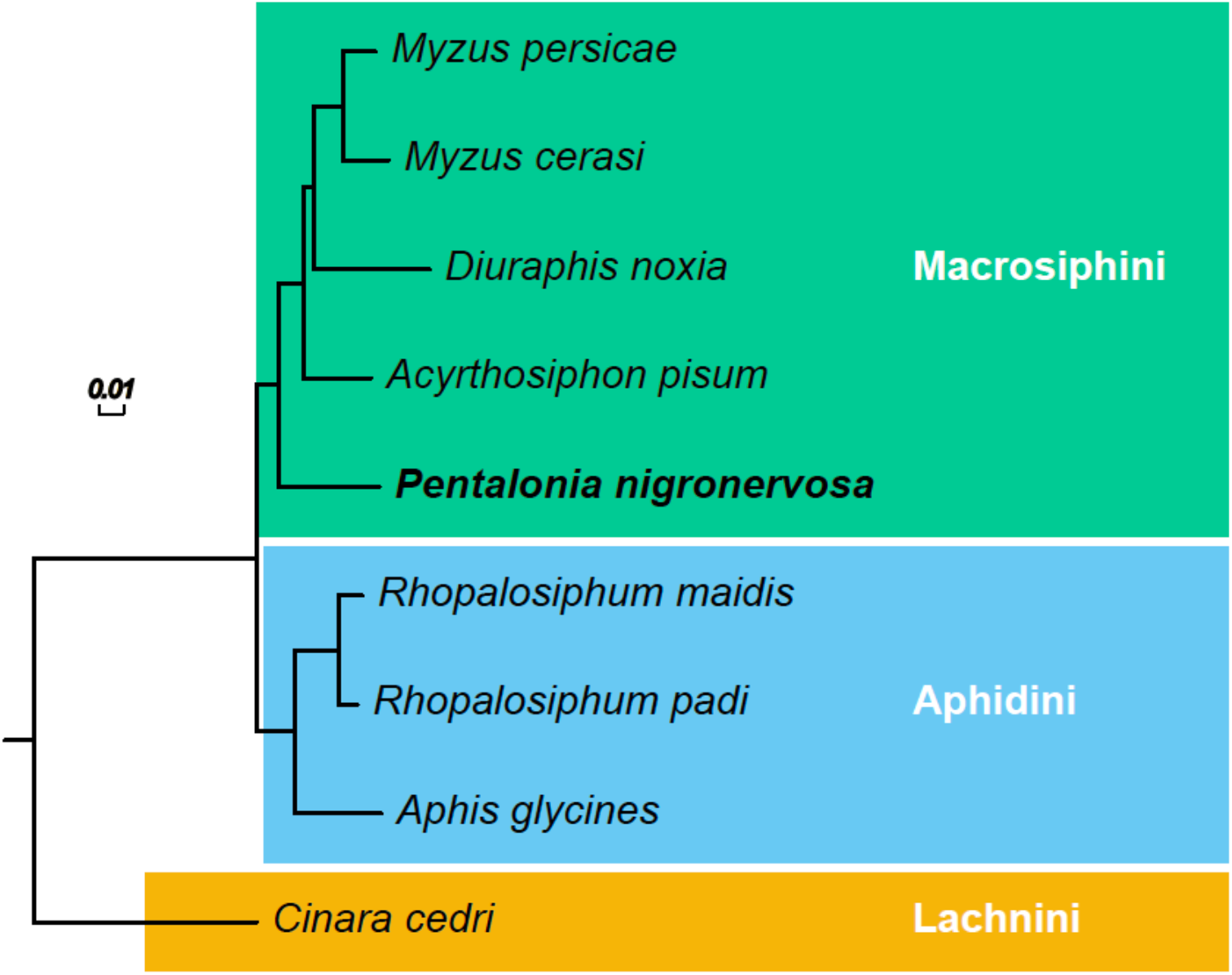
Maximum likelihood phylogeny of aphids based on a concatenated alignment of 4,721 conserved one-to-one orthologs. All branches received maximal support based on the Shimodaira-Hasegawa test (Shimodaira and Hasegawa 1999) implemented in FastTree (Price *et al.* 2009, 2010) with 1,000 resamples. Clades are coloured by aphid tribe. Branch lengths are in amino acid substitutions per site.

## Conclusions

Using a single Illumina short-read sequence library and high-coverage RNA-seq data we have generated a high-quality draft genome assembly and annotation of the banana aphid and simultaneously assembled the genomes of its *Buchnera* and *Wolbachia* symbiotic bacteria, providing an important genomic resource for the future study of this important pest. Furthermore, as an outgroup to other sequenced aphids from the tribe Macrosiphini, the banana aphid genome will enable more detailed comparative analysis of a group that includes a large proportion of the most damaging aphid crop pests (Van Emden and Harrington 2017) as well as important model species such as the pea aphid (Brisson and Stern 2006) and the green peach aphid (Mathers *et al.* 2017, 2020).

## Acknowledgements

TCM is funded by a BBSRC Future Leader Fellowship (BB/R01227X/1). The described work was supported by a CEPAMs grant (17.03.2) to SH, a Bill and Melinda Gates Foundation grant (OPP1087428) awarded to LT, the BBSRC Institute Strategy Program (BB/P012574/1) award to the John Innes Centre, and the John Innes Foundation. This research was supported in part by the NBI Computing Infrastructure for Science Group, which provides technical support and maintenance to the John Innes Centre’s high-performance computing cluster and storage systems.

