## Supplementary Figure for "Genome sequence of the banana aphid, *Pentalonia nigronervosa* Coquerel (Hemiptera: Aphididae) and its symbionts"

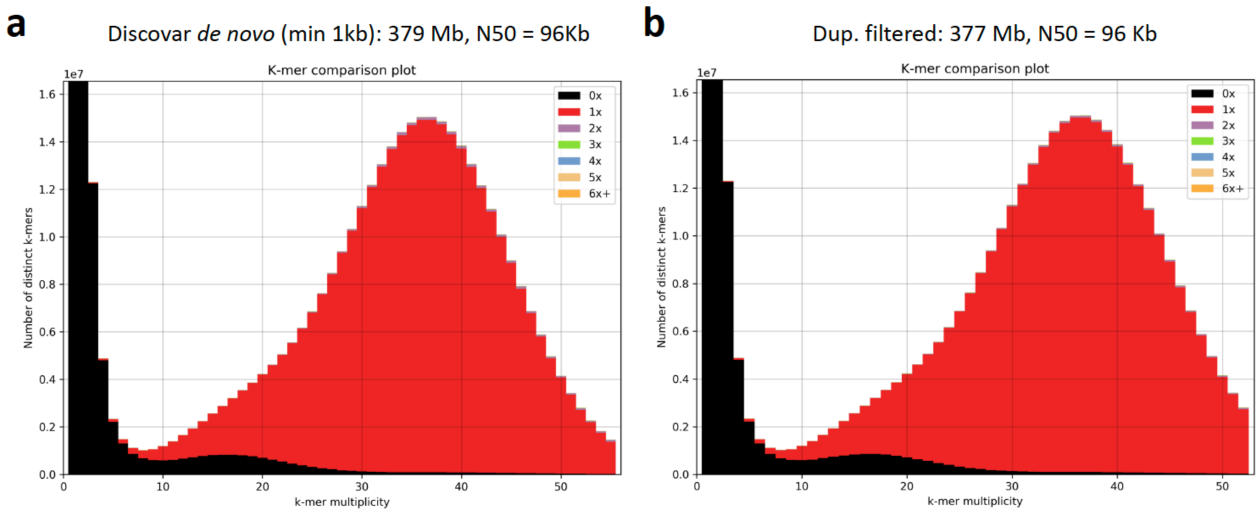


**Supplementary Figure 1:** KAT k-mer spectra plot comparing k-mer content of PCR-free *P. nigronervosa* Illumina reads to k-mer content of the initial discovar de novo genome assembly (**a**) and the same assembly after processing with our de-duplication pipeline (**b**) (see Methods). Colours indicate how many times fixed length words (k-mers) from the reads appear in the assembly. Red indicates k-mers found only once in the assembly, black indicates content present in the reads but missing from the assembly and other colours indicate k-mers that are duplicated in the assembly. The x-axis shows the number of times each k-mer is found in the reads (k-mer multiplicity) and the y-axis shows the count of distinct k-mers in 1x k-mer multiplicity bins.

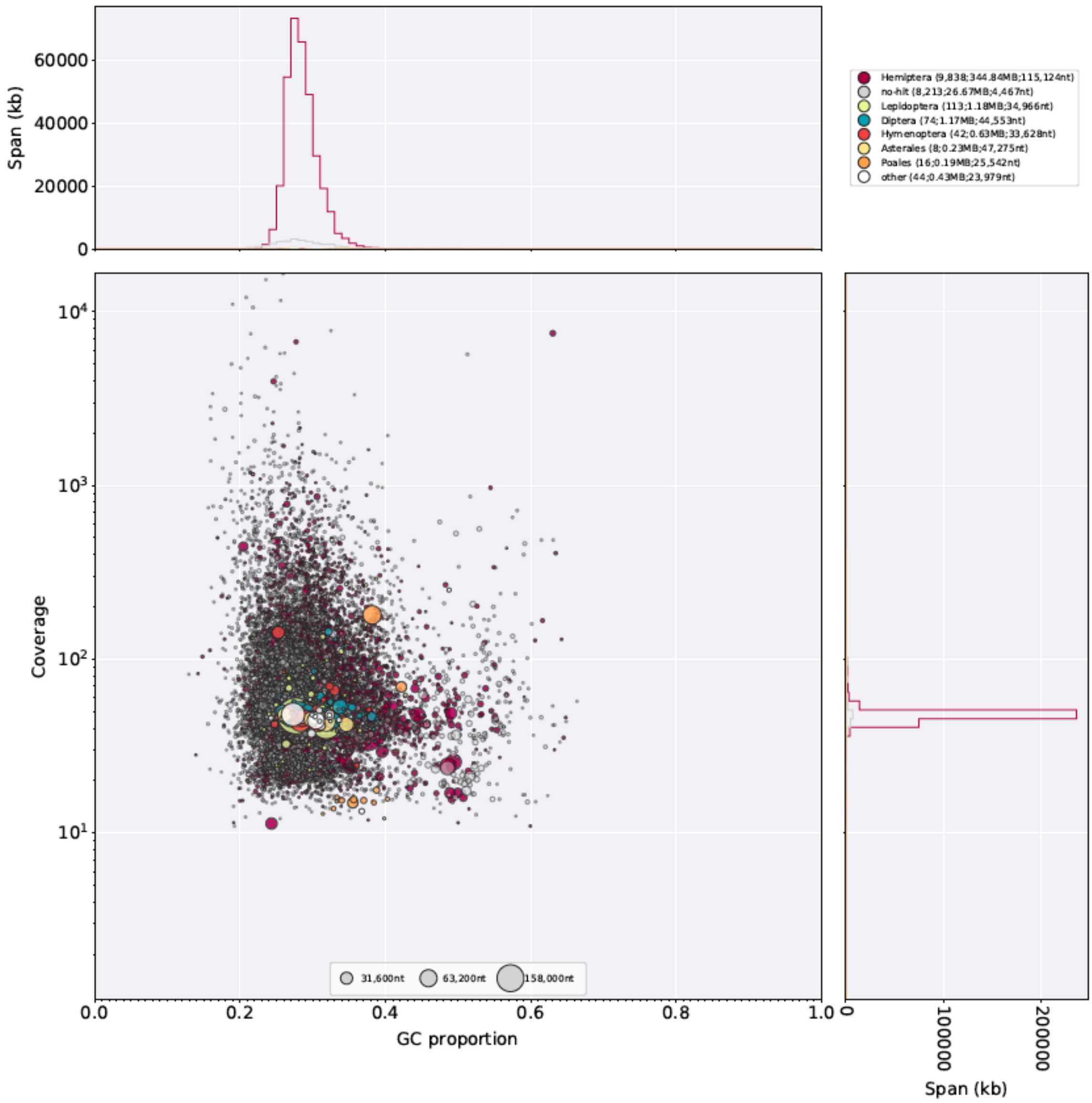


**Supplementary Figure 2:** Taxon-annotated GC content-coverage plot of the frozen *P. nigronervosa* genome assembly (Penig_v1). Each circle represents a scaffold in the assembly, scaled by length, and coloured by order-level NCBI taxonomy assigned by BlobTools. The X axis corresponds to the average GC content of each scaffold and the Y axis corresponds to the average coverage based on alignment of *P. nigronervosa* PCR-free Illumina short reads. Marginal histograms show cumulative genome content (in Kb) for bins of coverage (Y axis) and GC content (X axis).

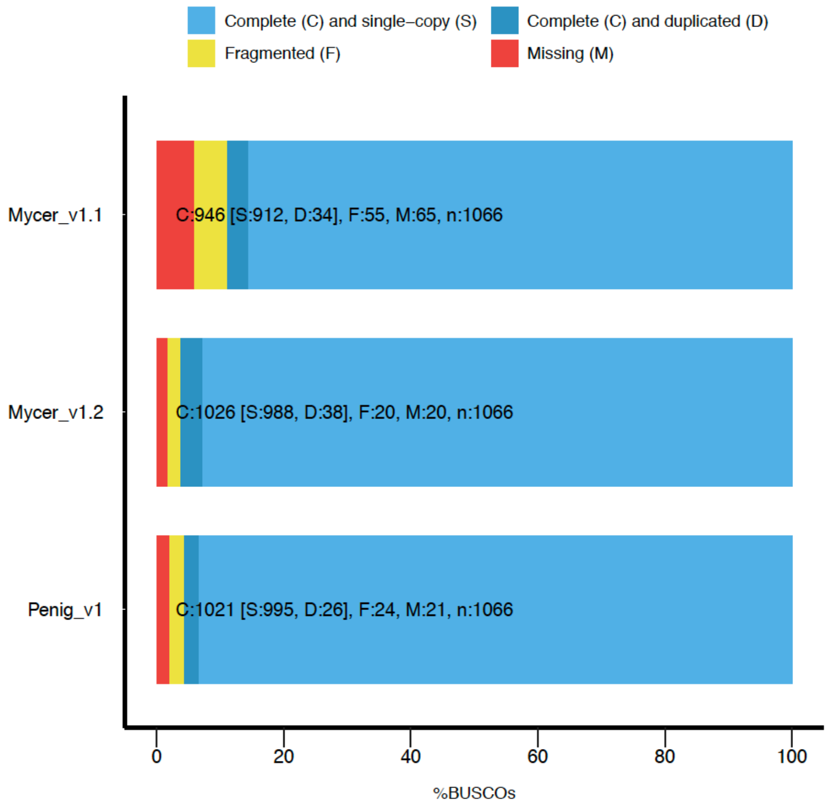


**Supplementary Figure 3:** BUSCO analysis of the *P. nigronervosa* v1 (Penig_v1), *M. cerasi* v1.1 (Mycer_v1.1) and *M. cerasi* v1.2 (Mycer_v1.2) gene sets (protein sequences). Where multiple transcripts of a gene where annotated we used the longest transcript to represent the gene. The protein sets were assessed using the Arthropoda gene set (n=1,066).

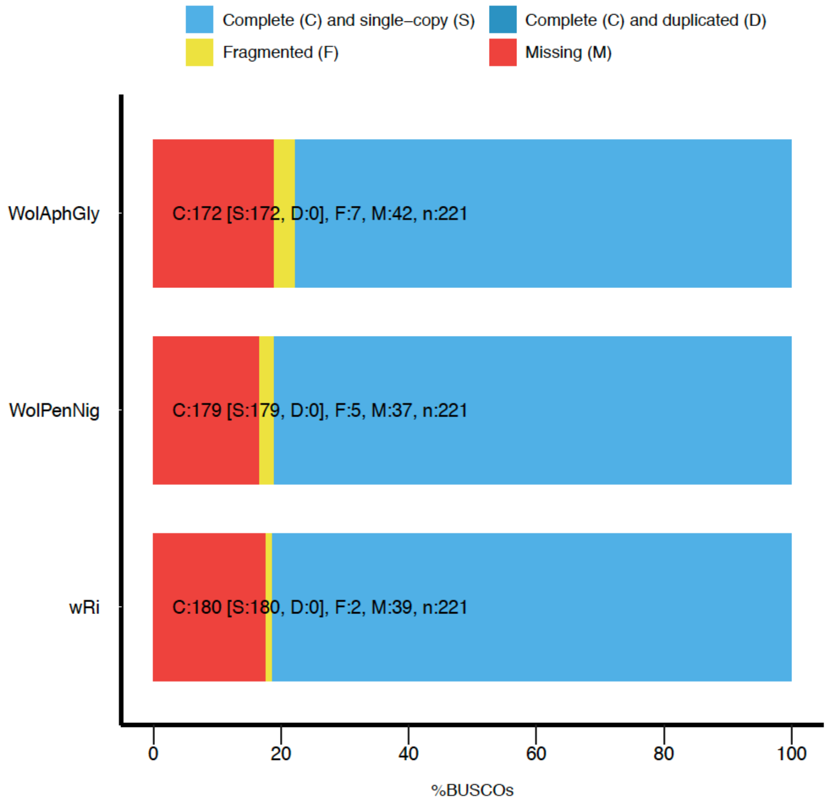


**Supplementary Figure 4:** The *P. nigronervosa* Wolbachia genome (WolPenNig) extracted from the discovar de novo genome assembly has similar completeness to the reference Wolbachia wRi genome (NCBI Reference Sequence: NC_012416.1) and a long-read assembly of a Wolbachia strain infecting the soybean aphid (WolAphGly). BUSCO gene-level completeness was assessed for each genome assembly using the proteobacteria gene set (n = 221).

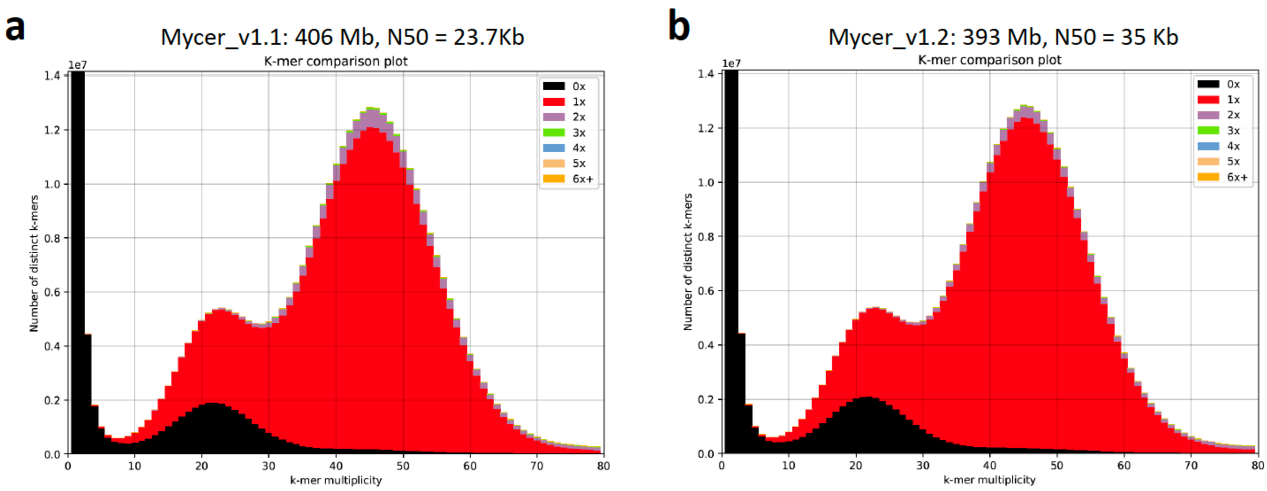


**Supplementary Figure 5:** KAT k-mer spectra plot comparing k-mer content of genomic PCR-free *Myzus cerasi* Illumina reads (NCBI accession: ERR2236145) to k-mer content of the published assembly of *M. cerasi* (**a**; Mycer_v1.1 [Thorpe et. al. 2018]) and the same assembly after processing with our de-duplication pipeline (**b**) (see Methods). The *stacked-histograms* are as described in **Supplementary Figure 1**.
